# *Enterococcus faecalis* V583 LuxS/AI-2 system is devoid of role in intra-species quorum-sensing but contributes to virulence in a *Drosophila* host model

**DOI:** 10.1101/344176

**Authors:** Frederic Gaspar, Neuza Teixeira, Natalia Montero, Tamara Aleksandrzak-Piekarczyk, Renata Matos, Bruno Gonzalez-Zorn, António Jacinto, Maria Teresa Barreto Crespo, Maria de Fátima Silva Lopes

**Affiliations:** iBET, Instituto de Biologia Experimental e Tecnológica, Apartado 12, 2781-901 Oeiras, Portugal.; Instituto de Tecnologia Química e Biológica António Xavier, Universidade Nova de Lisboa, Av. da República, 2780-157 Oeiras, Portugal.; Departamento de Sanidad Animal y VISAVET, Universidad Complutense de Madrid, Spain.; Institute of Biochemistry and Biophysics, Polish Academy of Sciences (IBB PAS), Pawińskiego 5a, 02-106 Warsaw, Poland.; CEDOC – Faculdade de Ciências Médicas, Universidade Nova de Lisboa, Lisboa, Portugal.

## Abstract

The AI-2 interspecies quorum-sensing molecule is produced by the LuxS enzyme and has been ascribed a role in virulence in several bacteria. The nosocomial pathogen *Enterococcus faecalis* inhabits several different environments where multispecies communities are established. However, despite the presence of a *luxS* gene in this pathogen, its role in *E. faecalis* pathogenesis has never been assessed. In the present work, we deleted the *luxS* gene from the vancomycin-resistant clinical isolate *E. faecalis* V583 and demonstrated the lack of AI-2 production by the mutant strain. Using microarrays and externally added (S)-4,5-dihydroxy-2,3-pentanedione we showed that AI-2 is not sensed by *E. faecalis* as a canonical quorum-sensing molecule and that the *luxS* mutation caused pleiotropic effects in gene expression, which could not be complemented by extracellularly added AI-2. These global differences in gene expression affected several gene functional roles, mainly those enrolled in metabolism and transport. Metabolic phenotyping of the *luxS* mutant, using Biolog plates, showed differences in utilization of galactose. AI-2 production by LuxS was shown to be irrelevant for some phenotypes related to the pathogenic potential of *E. faecalis* namely biofilm formation, adhesion to Caco-2 cells, resistance to oxidative stress and survival inside J-774 macrophages. However, the *luxS* mutant was attenuated when tested in the *Drosophila* septic injury model, as its deletion led to delayed fly death. Overall our findings show that differential gene expression related to the *luxS* mutation cannot be ascribed to quorum-sensing. Moreover, the role of LuxS appears to be limited to metabolism.

## Introduction

The species *Enterococcus faecalis* belongs to the normal microbiota of the GI tract of hosts as diverse as mammals and insects (Klein, 2003). They are also found in a variety of food products, namely milk and cheese produced in the south of Europe (Ogier & Serror, 2007). However, *E. faecalis* remains an important opportunistic pathogen and represents one of the main causes of nosocomial infections in the USA and Europe. Especially for immunocompromised patients, these infections include endocarditis, peritonitis, visceral abscesses, urinary infections or septicemia (Arias & Murray, 2012). Although *E. faecalis* is found in disparate environments, these are similar in the sense that they are all composed of multispecies communities. Recent years have been successful in the discovery of intra and inter-species communication, which is responsible for population density monitoring and for regulation of traits important for pathogenesis. In order to be successful inside the host, pathogenic bacteria produce virulence factors when they sense that it is worth to waste energy in their production. One environmental factor monitored by many pathogens is population density, either of its own population or of the population of a host’s endogenous flora (Parker & Sperandio, 2009). Intercellular communication is not the exception but, rather, the norm in the bacterial world (Shapiro, 2007). The process of sensing population density, called quorum-sensing (QS), is fundamental to coordinate certain behaviors of microbes. QS has been shown to regulate a variety of functions, including symbiosis, virulence, competence, motility, sporulation, mating, conjugation, antibiotic production, and biofilm formation (Pereira *et al*., 2013).

The small molecules that are produced, released and detected, and which mediate QS, are called autoinducers (Zhu & Pei, 2008). Most autoinducers are species-specific; however, one autoinducer, AI-2, and its synthase, LuxS, have been identified in many bacteria (Bassler, 1999; Surette *et al*., 1999; Sun *et al*., 2004; Pereira *et al*., 2009), and implicated in the regulation of many bacterial behaviors, including biofilm formation, competence, production of secondary metabolites like antibiotics, and virulence. While in some cases AI-2 is clearly acting through a canonical QS mechanism, in others a possibly primary and sometimes sole role in central metabolism has been proposed (Winzer *et al*., 2002a). The LuxS protein is an integral metabolic component of the activated methyl cycle (AMC). The AMC is a key metabolic pathway that generates S-adenosylmethionine (SAM) as an intermediate product. SAM bears a methyl group with a relatively high transfer potential, and is used by numerous methyltransferases to carry out cellular processes including nucleic acid and protein methylation, and detoxification of reactive metabolites. The product of the methyltransferase reaction is S-adenosylhomocysteine (SAH), and in the complete AMC, SAM is regenerated from SAH via homocysteine and methionine, ready for another round of methylation/transmethylation. The role of LuxS in the AMC is to catalyze the cleavage of SRH (S-ribosyl-L-homocysteine) to yield homocysteine and a by-product (S)-4,5-dihydroxy-2,3-pentanedione (DPD). DPD is the precursor of the family of related, inter-converting molecules collectively termed ‘AI-2’ (Schauder *et al*., 2001).

*E. faecalis* V583 strain was the first vancomycin-resistant clinical isolate reported in the United States (Sham *et al*., 1989) to be fully sequenced. Genomic database analysis has previously revealed (Surette *et al*., 1999; Paulsen *et al*., 2003) that this strain carries the gene encoding the putative LuxS protein, S-ribosylhomocysteine lyase (EC: 4.4.1.21), a 152 amino acid protein encoded by the *luxS gene* (*ef1182*) homologous to *luxS* from *V. harveyi*. Further *in vitro* analysis using either purified LuxS protein from *E. faecalis* overexpressed in *E. coli* strain BL21 (Schauder *et al*., 2001) or directly from *E. faecalis* V583 strain (Shao *et al*., 2012) revealed LuxS ability to produce AI-2. The *luxS* gene is disseminated among other recently sequenced *E. faecalis* genomes. In fact, a nucleotide BLAST (Altschul *et al*., 1997) with all the 55 finished and unfinished *E. faecalis* genomes and genome projects available in the microbial BLAST database from the National Center for Biotechnology Information (NCBI) revealed that the *ef1182* gene has 99% identity, or higher, in all the 55 hit results obtained.

Using a proteomic approach Shao *et al*. (2012) proposed that AI-2 affects biofilm formation and regulates some metabolic functions in *E. faecalis* V583 strain. However, the *luxS* role was not assessed. Considering the nature of ecological niches colonized by *E. faecalis*, it is important to first understand the role of LuxS/AI-2 system in *E. faecalis*, if and how *E. faecalis* senses the extracellular presence of AI-2 and if LuxS contributes to virulence in this species. We thus constructed a *luxS* deletion mutant in V583 and studied the effect of this mutation in several traits related to virulence, namely biofilm formation, adhesion to Caco-2 cells, survival inside macrophages, resistance to oxidative stress and *Drosophila* survival upon septic injury. Using microarrays and externally added DPD we also evaluated the ability of *E. faecalis* to sense AI-2 and the impact of the LuxS/AI-2 in gene expression.

## Materials and Methods

### Bacterial strains and general culture conditions

Bacterial strains used in this study are listed in Table 1. Enterococci were grown in M17 broth (BD, Franklin Lakes, NJ) supplemented with 0.5% (w/v) glucose (M17Glu) or M17Glu agar at 37°C, unless otherwise stated. *Escherichia coli* strains were grown in Luria–Bertani (LB) broth or on LB agar at 37°C. Antibiotics were used at the following concentrations: erythromycin, 30 μg ml^-1^ for *E. faecalis* and 150 μg ml^-1^ for *E. coli*; ampicillin, 80 μg ml^-1^.

**Table 1.**
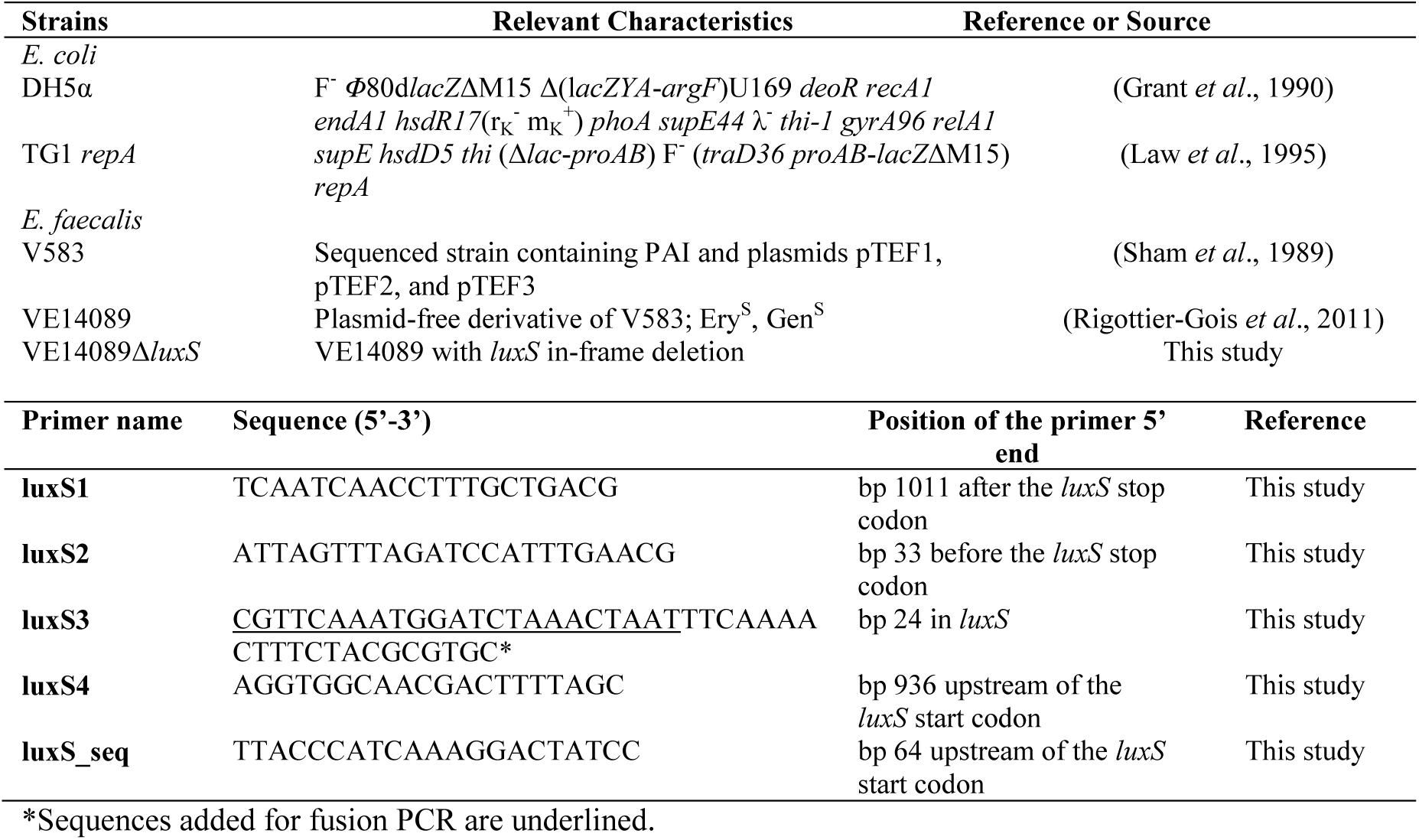
Strains and primers used in this study.

### General DNA techniques

General molecular biology techniques were performed by standard methods (Sambrook *et al*., 1989). Restriction enzymes, polymerases and T4 DNA ligase were used according to manufacturer’s instructions. PCR amplification was performed using a thermocycler (Biometra GmbH, Gottingen, Germany). When necessary, PCR products and DNA restriction fragments were purified with QIAquick purification kits (Qiagen, Hilden, Germany). Plasmids were purified using the QIAprep Spin Miniprep kit (Qiagen, Hilden, Germany). Electrotransformation of *E. coli* and *E. faecalis* was carried out as described by Dower *et al*. (1988) and Dunny *et al*. (1991), using a Gene Pulser apparatus (Bio-Rad Laboratories, Hercules, CA). Genomic DNA fragments and plasmid inserts were sequenced at Baseclear (Netherlands).

### Construction of in-frame *luxS* deletion mutant in strain VE14089

We studied the activity of LuxS in the strain VE14089 (Rigottier-Gois *et al*., 2011), which is a V583 derivative cured of its plasmids, and referred to as WT hereafter, a genetically tractable strain compared to the original V583 (Matos *et al*., 2013). Besides removing the effect of plasmid-encoded genes and making easier the markerless in-frame *luxS* deletion mutagenesis, using strain VE14089 allowed us to test the strain in mammalian cell culture because it is gentamicin sensitive, as opposed to the parental V583. The markerless *luxS* deletion mutant of *E. faecalis* strain VE14089 was constructed essentially as described by Gaspar *et al*. (2009). Briefly, 5’ and 3’ flanking regions of luxS were amplified from chromosomal DNA of each strain by PCR with primers luxS1, luxS2, luxS3, and luxS4 (Table 1). The two cognate PCR fragments were fused by PCR using the external primers luxS1 and luxS4, and the resulting product was cloned into pGEM-T (Promega Corporation, Fitchburg, WI). The inserted PCR fragment was removed from its cloning vector by restriction enzymes and subsequently cloned into pG+host9 plasmid (Maguin *et al*., 1996), which was then electroporated into *E. faecalis*. The *luxS* single- and double-crossover mutants were selected as described by Brinster *et al*. (2007). Successful targeted mutations of *luxS* in strains VE14089 were first identified by PCR screening and then confirmed by Southern blot analysis.

### Quantification of AI-2

To monitor extracellular AI-2 activity in cell cultures during growth, cell-free culture supernatants were prepared by filtration and then analyzed for AI-2 activity. AI-2 quantification was done using a LuxP-FRET-based reporter (FRET — fluorescence resonance energy transfer), as established by Rajmani *et al*. (2007) and optimized for 96-well plate reading by Marques *et al*. (2011). The binding of AI-2 to the CFP-LuxP-YFP chimeric protein causes a dose-dependent decrease in the FRET signal, and concentration can be determined by comparing the FRET ratios (527 nm/485 nm) of each sample with a calibration curve performed with AI-2 samples of known concentration. Concentrations between 1 and 60 μM DPD (Omm Scientific, Inc., Dallas, TX) were used for the calibration curve, corresponding to the linear range of this assay. All assays were performed in duplicate.

### Microarray Analysis

For all experiments, the strains were inoculated in 5 mL 2×YTGlu broth and grown for 16 h without shaking at 37 °C. The culture was then diluted in pre-warmed to 37°C 2×YT broth, adjusting the bacterial suspension density to the 1.0 McFarland standard. This pre-culture was diluted 1:100 (v/v) in 50 mL pre-warmed to 37°C 2×YT broth, and incubated, with 150 rpm agitation in an orbital shaker (Innova, Edison, New Jersey), at 37°C, until, for each condition studied, samples for RNA isolation were collected and processed accordingly. When needed, 10 μM DPD was added 15 min prior to the RNA collection. The whole process was performed twice, independently. Immediate RNA sample stabilization and protection was achieved using the RNAprotect Bacteria Reagent (Qiagen, Hilden, Germany). Total RNA purification was performed using the RNeasy Midi Kit (Qiagen, Hilden, Germany), DNA digestions were executed after RNA isolation using DNase I recombinant, RNase-free (Roche Applied Science, Penzberg, Germany) and repeated when necessary, and final RNA clean up and concentration was carried out with RNeasy MinElute Cleanup Kit (Qiagen, Hilden, Germany), all according to the manufacturer’s instructions. Integrity and overall quality of the total RNA preparations, but also DNA contamination, were evaluated by native agarose gel electrophoresis and by PCR, respectively, and the corresponding RNA concentrations were measured using a NanoDrop spectrophotometer (NanoDrop Technologies, Inc., Thermo Fisher Scientific Inc., Wilmington, DE). Microarray comparative genomic hybridization analysis was carried out using the microarray analysis platform of NimbleGen Technologies (Roche NimbleGen, Madison, WI). The chosen expression microarray, 4×72k format (Catalogue Number: A7980-00-01, Design Name: 080625 Efae V583 EXP X4), covers 3114 genes represented with 11 60mer probes per gene and 2 replicates per probe. cDNA synthesis, labelling, hybridization, and data acquisition were performed by NimbleGen (Reykjavik, Iceland). Image analysis was performed with the NimbleScan software (v2.6) (Roche NimbleGen Inc., Madison, WI), and feature intensities were exported as pair files, after background correction and quantile normalization. All data analysis was carried out using the ArrayStar 3.0 software package (DNAStar, Madison, WI). Robust multichip averaging (RMA) algorithm and quantile normalization were used for probe summarization and normalization and applied to the entire data set, which consisted of two biological replicates for each condition. Statistical analyses were carried out with the normalized data using a moderated t-test with false discovery rate (FDR) multiple-test correction (Benjamini-Hochberg) to determine differential transcript abundance. Changes in transcript abundance were considered significant if they met the following criteria: P value < 0.05 and | log2-ratio | > 5.0. Results can be accessed through GEO accession number GSE52374.

### Global phenotypic testing of carbon source utilization

*E. faecalis* ability to grow from different carbon sources was measured globally in duplicate by the Phenotype MicroArrays system (Biolog, USA) according to manufacturer’s instructions. Briefly, *E. faecalis* strains (VE14089 strain and its isogenic *luxS* mutant) were streaked on plates containing M17Glu agar. Colonies were scraped from the plates and suspended in IF-0a inoculating fluid (Biolog) with growth supplements and Biolog redox dye mixture according to standard protocols recommended by Biolog for *Enterococcus* species. 100 µl aliquots were added to each well of carbon source plates (PM1 and PM2). The PM1 and PM2 B i ol og assays assess the ability of a bacterium to utilize any of 190 carbon compounds used as the sole carbon source. The plates were incubated at 37°C in aerobic conditions in the OmniLog incubator plate reader, and cellular respiration was measured kinetically by determining the colorimetric reduction of a tetrazolium dye. Data were collected approximately every 10 min over a 48-h period. Data were analyzed with the Biolog Kinetic and Parametric software.

### Biofilm assay on polystyrene microtiter plates

Biofilm formation on polystyrene was quantified with crystal violet staining method as previously described (Thomas *et al*., 2009). Briefly, strains were grown for 16 h in 2×YT (BD, Franklin Lakes, NJ) supplemented with 0.5% glucose (2×YTGlu) broth, at 37°C, and the culture was subsequently diluted 1:100 (v/v) in pre-warmed to 37°C 2×YT broth. 200 μl of the diluted cell suspension was used to inoculate sterile 96-well polystyrene microtiter plates (Sarstedt, Nümbrecht, Germany). Biofilms were processed after 24 h incubation at 37°C, as described above. Each assay was performed in hexuplicate, repeated twice, and the overall significance of the differences was determined by a two-tailed unpaired t-test. All experiments included a blank well (medium without any inoculum).

### Adherence assay

The ability of *E. faecalis* strains VE14089 and VE14089*ΔluxS* to adhere to Caco-2 cells (obtained from the cell bank of the Centro de Investigaciones Biologicas CIB-CSIC, Madrid) was determined as previously described (Olier *et al*., 2003), with minor modifications. On 24-well tissue-culture plates, a 95% confluent monolayer of Caco-2 cells was infected with an *E. faecalis* bacterial suspension, with a corresponding multiplicity of infection (MOI) of ~50. Adhesion of *E. faecalis* cells to Caco-2 cells was allowed to occur for 2 h at 37°C. They were then washed 3 times with PBS. Adherent bacteria were harvested after lysis of the cell monolayers with Triton X-100 and suitable dilutions of the lysates were plated. The plates were subsequently incubated for 24-48 h at 37°C and CFU values for viable bacteria were determined. Adherence assays were done in triplicate, and the overall significance of the differences was determined by a two-tailed unpaired t-test.

### H_2_O_2_ challenging conditions

H_2_O_2_ challenge was performed based on the method described by Giard *et al*. (2006), with some adaptations. Briefly, WT and mutant cells were inoculated in M17Glu broth and grown for 16 h without shaking at 37°C. The culture was then diluted 1:100 (v/v) in pre-warmed to 37°C M17Glu broth and incubated, with 150 rpm agitation in an orbital shaker (Innova, Edison, New Jersey), at 37 °C, until reaching an OD at 600 nm of approximately 0.5. The cells were harvested by centrifugation and resuspended in M17 broth with 7 mM H_2_O_2._ These cultures were placed into a 37°C water bath and, every 2 h for 6 h, samples were taken and plated in M17Glu agar. The number of CFU was determined after incubation at 37°C. The growth of the mutant and WT cells in the absence of peroxide stress was previously determined and did not reveal any difference. Each point is the mean of four independent experiments, each with duplicate plating, and the statistical comparison of means was performed using a two-tailed unpaired t-test. Survival at any given time point was determined as the ratio of the number of CFU after treatment to the number of CFU at the zero time point.

### Macrophage Survival Assay

The macrophage survival assay was mainly performed as described by Bennett *et al*. (2007) with some modifications. Confluent J774.A1 (mouse monocytes - macrophages) monolayers were infected with an overnight bacterial culture. Approximately 4 × 10^6^ bacteria were added to J774.A1 monolayers, to yield a MOI of approximately 10, and were incubated at 37°C in 5% CO_2_ atmosphere for 1 h to allow bacterial adherence and entry, after which gentamicin (250 μg ml^-1^) was added to the cultures to kill extracellular bacteria. At various time points after infection, 1% Triton X-100 in PBS was used to lyse cells. Lysates were then serially diluted and inoculated on Brain heart infusion (BHI) (Oxoid Ltd, Basingstoke, England) plates to enumerate viable intracellular bacteria. The assays were performed 5 times and results are reported as intracellular Survival Index (SI), i.e. the per cent (mean) of the internalized CFUs at each analyzed time post-infection that survived after phagocytosis.

### *Drosophila* infection with *E. faecalis*

*Drosophila* infection was performed according to Teixeira *et al.* (2013). Oregon R male flies were injected with 50 nl of bacteria at OD600 0.02 from one of the strains, VE14089 and VE14089*ΔluxS*. As control, flies were injected with the same volume of BHI medium. Male flies were anesthetized with CO_2_ and the injections were carried out with a pulled glass capillary needle using a nanoinjector (Nanoliter 2000, World Precision Instruments). Injected flies were placed at 29°C, 65% humidity. Twenty-five flies were assayed for each survival curve. Each experiment was repeated three times, making a total of 75 flies tested per strain. Death was recorded at 0, 4, 6, 8, 10, 12, 14 and 24h hours post-injection. Statistical analysis of *Drosophila* survival was performed using GraphPad Prism software version 5.03. Survival curves were compared using Log-rank and Gehan-Breslow-Wilcoxon tests.

## Results

### *luxS* mutagenesis depletes AI-2 activity

In order to assess LuxS activity in VE14089 strain, we measured the ability of this bacterium to produce AI-2. Produced AI-2 concentrations were deduced from the calibration curve shown in Figure 1. As shown in Fig. 2A, cell-free culture supernatants of the VE14089 strain induced a growth phase dependent signal, above the detection method threshold, indicating that AI-2 molecules were produced by the VE14089 strain. In contrast to VE14089 wild-type strain, no AI-2 activity was detectable in the supernatant of the *luxS* mutant, regardless of cell density (Fig. 2B). The *luxS* deletion abolished the production of AI-2, demonstrating that LuxS is the key determinant in the AI-2 production process in VE14089.

**Figure 1:**
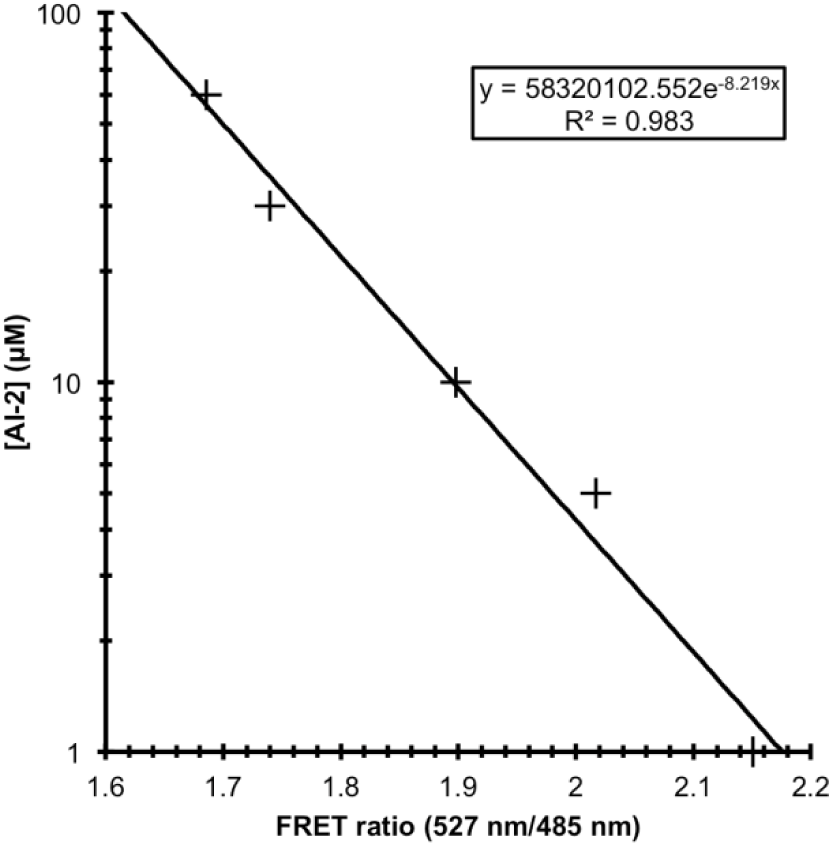
Calibration curve performed with AI-2 samples of known concentration. Concentrations of 1, 5, 10, 30 and 60 μM AI-2 were used for the calibration curve, corresponding to the linear range of this assay. Binding of AI-2 to the CFP-LuxP-Y FP fusion protein causes a dose-dependent decrease in the FRET signal, and concentration can be determined by comparing the FRET ratios (527 nm/485 nm) of each sample with the calibration curve.

### Exogenously added AI-2 does not complement *luxS* deletion

We were also interested in discriminating between the role of *luxS* and the ability of *E. faecalis* to sense AI-2 as a quorum-sensing molecule. We thus performed a transcriptomic analysis of VE14089, VE14089*ΔluxS* and VE14089*ΔluxS* supplemented with DPD. From previous experiments (Fig. 2), we knew that high AI-2 levels were achieved in the transition between late-exponential and early-stationary phase. Thus, cells were collected at late-exponential phase, which guarantees that, for the tested conditions, AI-2 is present extracellularly for the parental strain and that a possible effect for its presence may be monitored. The *luxS* mutant supplementation with DPD was performed 15 min prior to the cell harvesting, which did not notably interfere with VE14089Δ*luxS* growth, and came close to the level observed for the parental strain (Fig. S1). We chose to use synthetic AI-2 molecules in the form of DPD, as previously done in other studies (Ahmed *et al*., 2009; Kint *et al*., 2009; Armbruster & Swords, 2010), to avoid using purified culture filtrates that might have been misleading, as they contain complex mixture of other signals to which bacteria could respond (Vendeville *et al*., 2005). Genes regulated by AI-2 were identified by performing a pair-wise comparison of gene expression in VE14089Δ*luxS* and VE14089Δ*luxS* supplemented with 10 μM DPD. Pair-wise comparisons of gene transcription in VE14089 and VE14089Δ*luxS*, supplemented or not with 10 μM DPD, allowed us to determine changes in gene expression affected uniquely by the *luxS* mutation or by the *luxS* mutation along with the lack of extracellular *in vitro* presence of AI-2, respectively.

**Figure 2.**
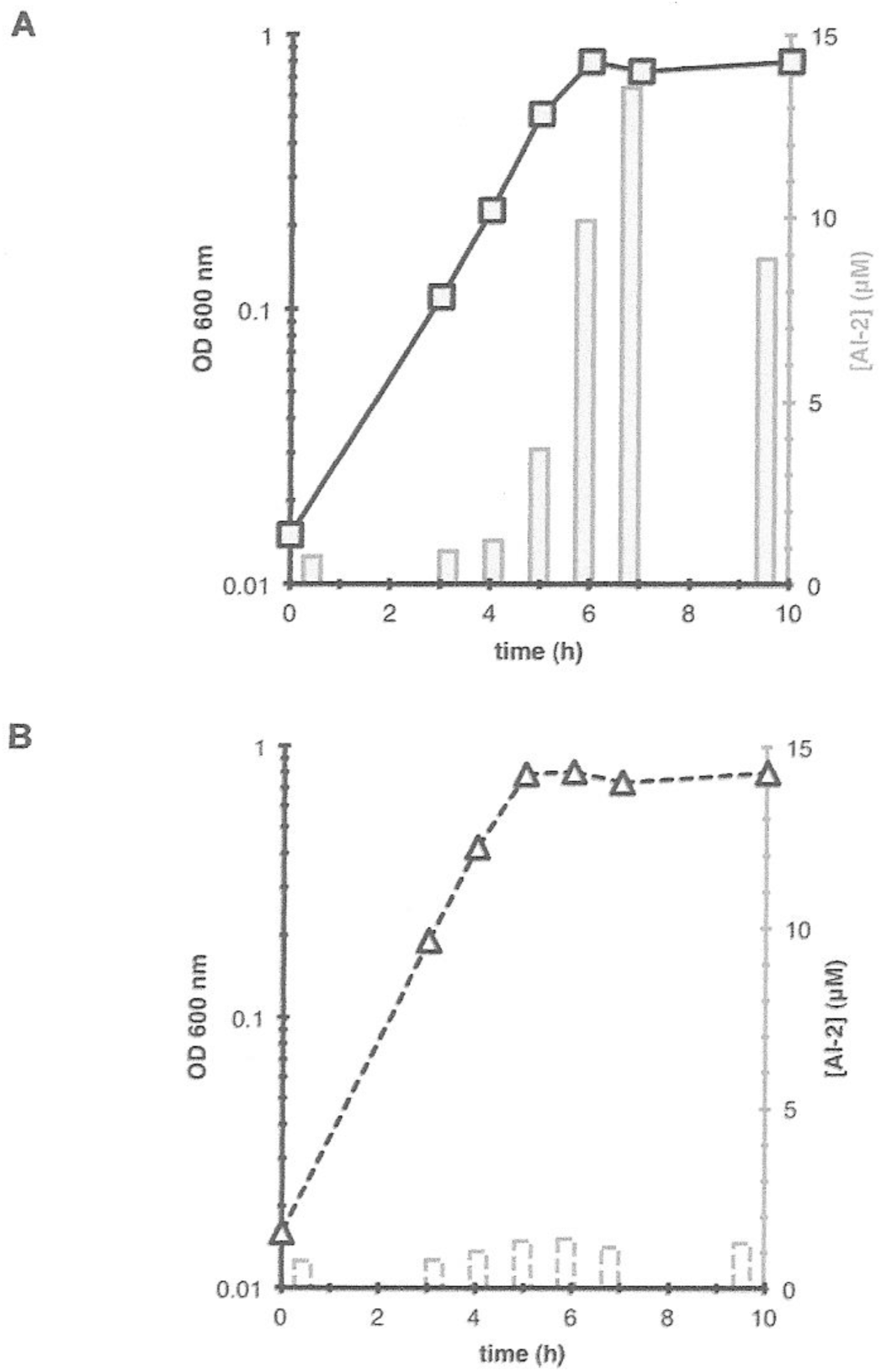
*E. faecalis* growth curves and AI-2 production. (A) VE14089; (B) VE14089*ΔluxS*. Strains were grown in 2xYT medium. During growth, monitored by measuring OD at 600 nm, culture samples were taken and AI-2 concentration in the supernatant was measured using the calibration curve from Figure 1.

Fundamental changes in gene expression were observed when comparing the wild-type strain and *luxS* mutant, affecting a total of 113 genes of all 3114 chromosomal genes present in the microarray, corresponding to 3.6% of the whole genome and considering fold-change values above 5 (Table S1). The full 113 differentially regulated genes in the *luxS* mutant, 70 upregulated and 43 downregulated, were found to be uniquely affected by the *luxS* mutation, independently of DPD addition, when compared to the parental strain. In fact, transcription patterns remained unaffected by addition of DPD to VE14089Δ*luxS*, when compared to VE14089Δ*luxS* (Table S1), indicating the absence of genes that could be responding to the signaling molecule AI-2.

### *luxS* deletion affects expression of genes mainly involved in energy metabolism, signal transduction and transport and binding

The *luxS* mutation resulted in changes in nearly every cellular process (Fig. 3), and the affected genes were distributed throughout the genome (Table S1). Unknown function or hypothetical protein genes contributed the largest fraction, more than 35%, followed by genes required for transport and binding, signal transduction, energy metabolism, cell envelope, and regulatory functions.

**Figure 3.**
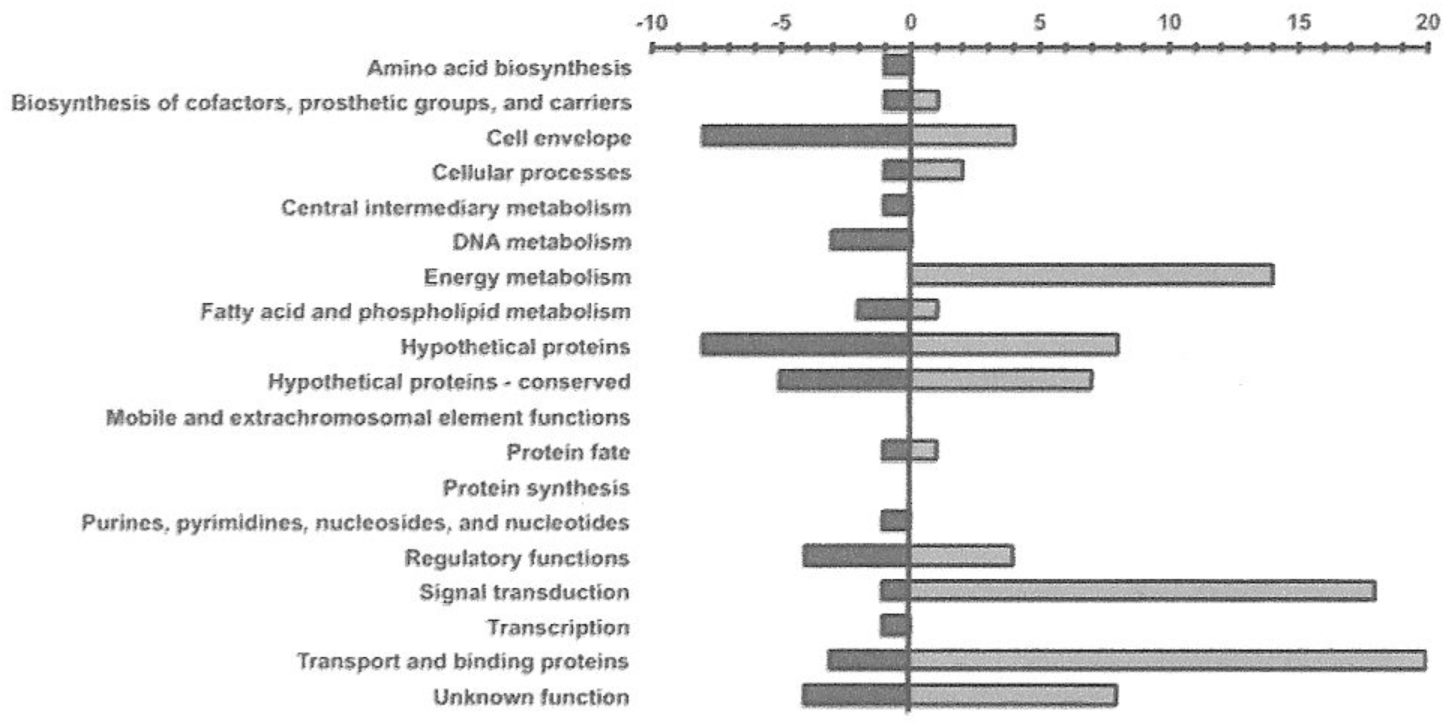
Number and role of differentially expressed genes associated to *luxS* mutation. Genes found to be affected by the *luxS* mutation, independent of (S)-4,5-dihydroxy-2,3-pentanedione addition, when compared to the parental strain VE14089, are grouped according to JCVI (http://cmr.jcvi.org) cellular main role. Differentially expressed genes with | log2-ratios | > 5.0 are presented (up-regulated,in light grey and shown as positive; downregulated,in dark grey and shown as negative). The genes with more than one cellular main role were counted twice.

In our study, most of the observed up-regulated genes in the *luxS* mutant, which also had fold-change values above 10, had functions related with energy metabolism (Table 2). All differentially expressed components of the phosphoenolpyruvate (PEP) transport system (PTS) were strongly upregulated, between 5 and 75-fold. Overall, when compared to VE14089 strain, the *luxS* mutant displays an increased transcription of genes involved in the transport and utilization of less preferred carbon sources, including mannose, cellobiose, mannitol, fructose, sorbitol/glucitol and gluconate.

**Table 2.**
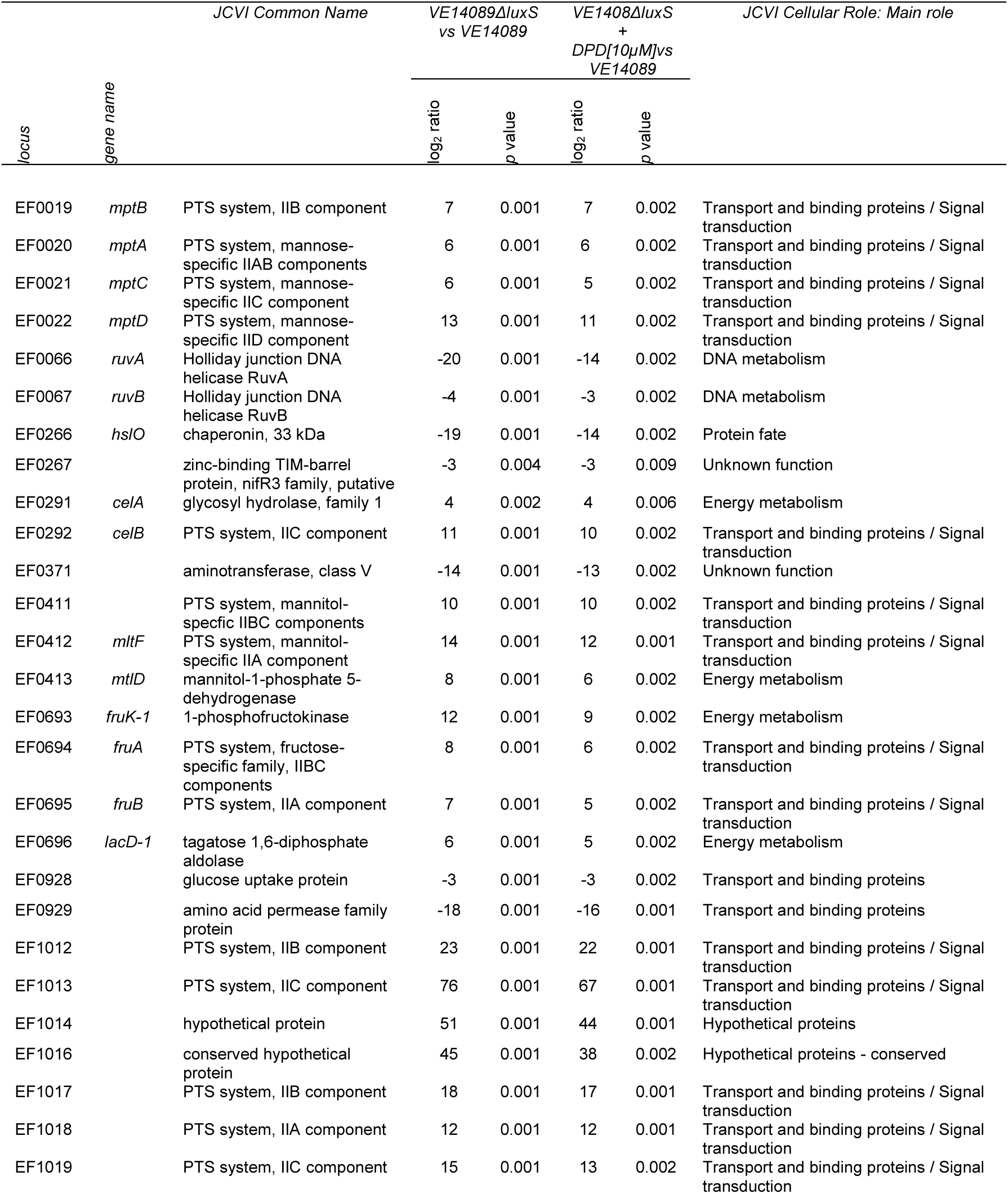
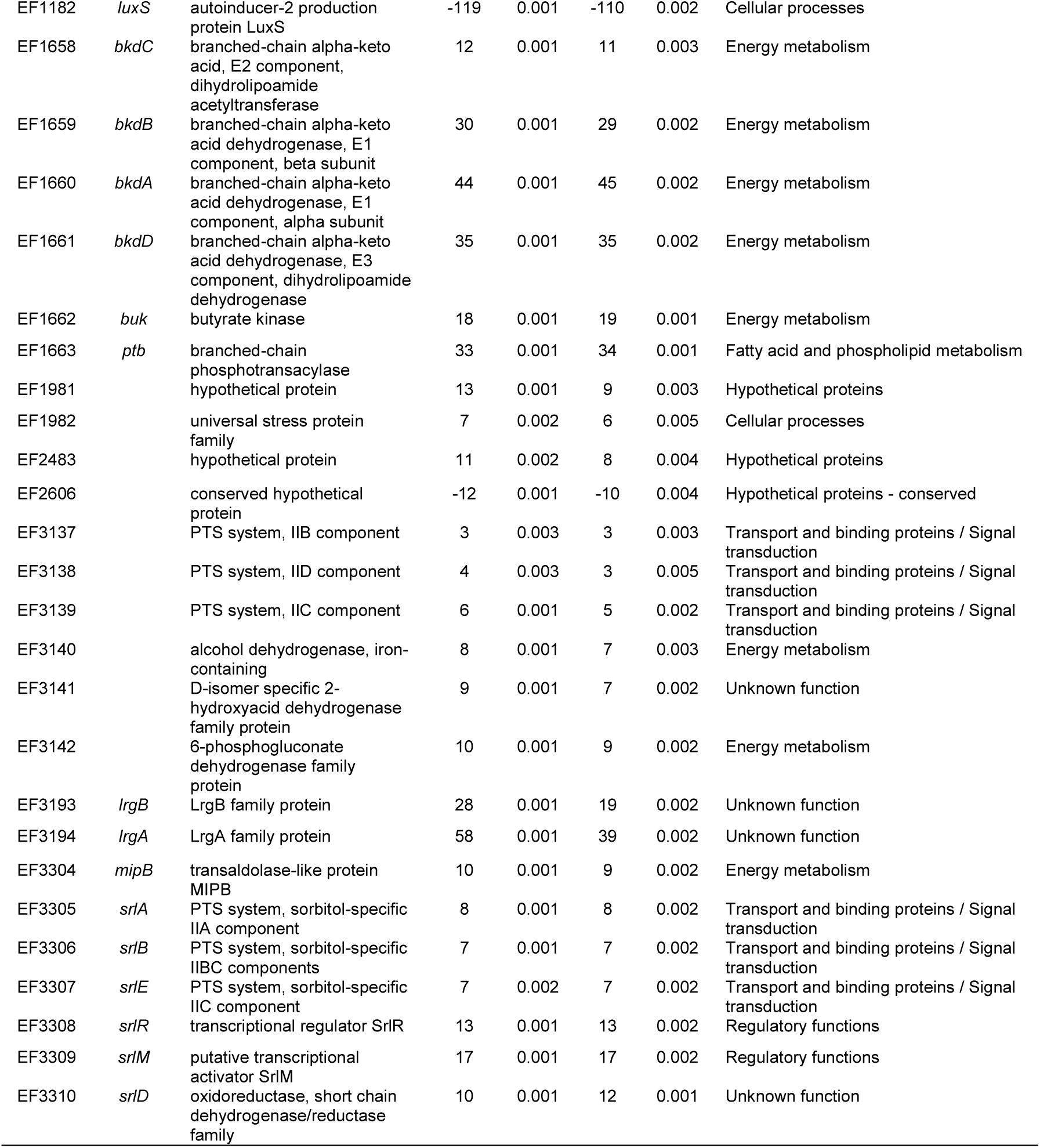
List of differentially regulated genes with fold-change values above 10, in at least one of the pairwise comparisons: VE14089 and VE14089*ΔluxS*, and VE14089 and VE14089*ΔluxS* supplemented with 10 μM DPD. Genes belonging to the same transcriptional unit, not significantly expressed, or with a fold change below 10 are also indicated.

In particular, in the *luxS* mutant, the *E. faecalis* branched-chain alpha-keto acid dehydrogenase (BCKDH) complex, encoded by the gene cluster *ptb-buk-bkdDABC* (*ef1663-ef1658*), was strongly up-regulated (Table 2), indicating an increased usage of branched-chain amino acid (BCAA). Altogether, the *luxS* mutant presents increased carbon flux from sources other than hexose sugars that are rapidly depleted from 2×YT growth media, to which no glucose was added.

The extensive transcriptional changes in the *luxS* mutant lead us to ask if there were any regulatory networks putatively regulated by pleiotropic regulators. Using Virtual Footprint v3.0, and allowing 1 sequence mismatch, we searched the *E. faecalis* V583 genome for the catabolite responsive elements (cre), using query consensus sequence WTGWAARCGYWWWCW, developed for *E. faecalis* (Opsata *et al*., 2010), the previously described sequence of a rex-box, TGTGANNNNNNTCACA (Mehmeti *et al*., 2011), and the sigma-54 consensus sequence YTGGCACNNNNNTTGCW (Opsata *et al*., 2010), developed for *B. subtilis*. Several promoter regions of differentially expressed genes in the *luxS* mutant were identified (Table 3), suggesting that *E. faecalis* responds and adapts to *luxS* deletion through a highly regulated and coordinated response.in which catabolite repression plays an important role. As our transcriptomic data predicted an overall change in metabolic profile due to absence of the LuxS protein, we performed a Biolog phenotypic array using plates covering 190 different carbon sources. Among them, the presence of one carbon source, D-galactose, led to clear and statistically significant differences in metabolic activity between the two strains. The *luxS* mutant was faster (Lag phase of the mutant strain was 8 h and that of the wt strain was 20 h, p<0.05) in using D-galactose, showing also nearly the double growth when compared to the wild-type strain (14561 vs 26770 of area under growth, for the wild-type and mutant, respectively, p<0.05). Although we cannot establish a correlation between the Biolog results and those of the transcriptomic analysis, due to different experimental conditions intrinsic to the experimental setup, both assays allowed for detection of differences in metabolism between the wild-type *E. faecalis* and the *luxS* mutant.

**Table 3.** Differential expression of genes associated with the interaction with transcriptional regulators, with the consensus motif localized in the intergenic region (int) or in the coding region (cod) in the VE14089Δ*luxS* mutant when compared to VE14089, independently of extracellular added DPD. DOWN: down-regulation (*P*-value < 0.05, log_2_ ratio < −10), UP: up-regulation (*P*-value < 0.05, log_2_ ratio > 10).

| | Gene name | $\log_2$ ratio | Motif (localization) |
| --- | --- | --- | --- |
| **response to carbon usage:** |  |  |  |
| fructose usage | *ef00693-6* | UP | cre (int) |
| cellobiose usage | *ef1012* | UP | cre (int) |
| cellobiose usage | *ef1013?* | UP | cre (int) |
| *ptb-buk-bkdDABC* | *ef1663-58* | UP | cre (int) |
| BCAA usage |  |  |  |
| *ptb-buk-bkdDABC* | *ef1663-58* | UP | cre (cod) |
| BCAA usage |  |  |  |
| gluconate usage | *ef3142-37* | UP | cre (int) |
| amino acid permease | *ef0929* | DOWN | cre (cod) |
| **response to NADH/NAD<sup>+</sup> ratio:** |  |  |  |
| *mptBACD ef0019-22* | *ef0020* | UP | rex (int) |
| **response to the interaction with the environment:** |  |  |  |
| *mptBACD* | *ef0019-22* | UP | sigma-54 (int) |
| cellobiose usage PTS | *ef1012* | UP | sigma-54 (int) |
| cellobiose usage PTS | *ef1017-8* | UP | sigma-54 (int) |
| *murAA* / HP peptidoglycan biosynthesis | *ef2605-4* | DOWN | sigma-54 (cod) |
| gluconate usage (in *ef3142-37* operon) | *ef3139* | UP | sigma-54 (cod) |

### *luxS* does not affect traits known to be relevant to *E. faecalis* virulence

Besides the downregulation of some genes of the *epa* cluster (Table S1), namely *ef2197*, none of the other potential virulence factors described by Manson & Gilmore (2006) were significantly regulated in the *luxS* mutant. These findings suggest that LuxS/AI-2 system may not have implications for *E. faecalis* virulence. In order to confirm this, we characterized the effect of the *luxS* deletion in VE14089 regarding traits that may be involved in the host-pathogen relationship, such as host recognition, adhesion and survival. For this purpose, four phenotypic experiments were carried out: biofilm formation on an abiotic surface, adherence to Caco-2 cells, resistance to oxidative stress, and survival inside macrophages. When using a two-tailed unpaired t-test, no significant differences were observed between VE14089 and VE14089Δ*luxS* strains regarding biofilm formation (Fig. 4A), adhesion to Caco-2 cells (Fig. 4B), H_2_O_2_ challenge (Fig. 4C), survival inside macrophages (Fig. 4D) and uptake by macrophages (8.2% uptake (± 2.5) at MOI 12.2 (± 2.7) by VE14089; 7.7% (± 2.2) uptake at MOI 14.3 (± 2.6) by VE14089Δ*luxS*)). Also, the *luxS* deletion did not affect growth of the mutant, having no obvious difference in growth from mid-exponential onwards and reaching stationary phase at the same time (Fig. S1). All these results indicate that, at least in the conditions used here, *E. faecalis* LuxS does not seem to be involved in the regulation of growth, biofilm formation, adherence to epithelial cells, resistance to oxidative stress and survival inside macrophages.

**Figure 4.**
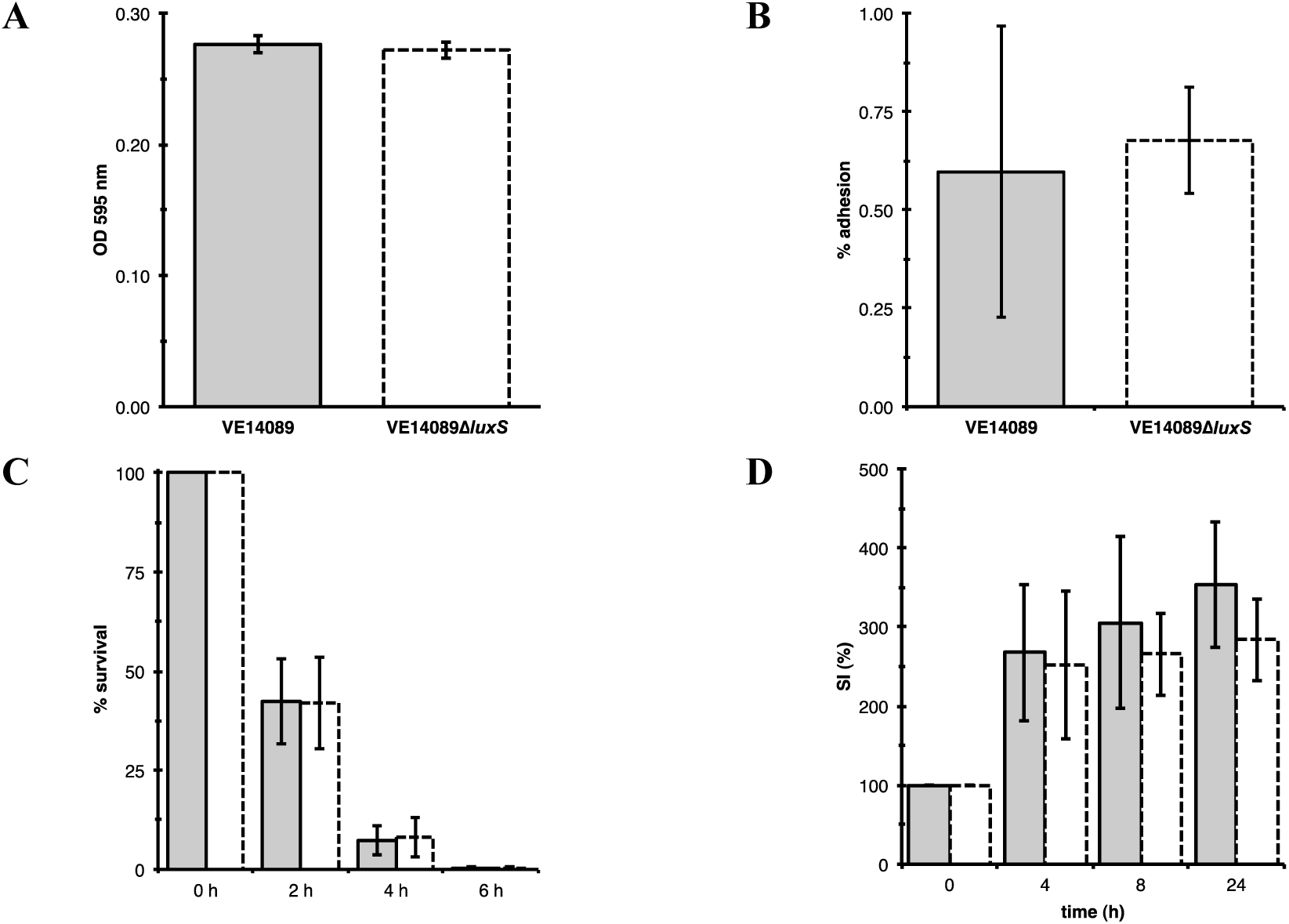
Phenotypic characterization of the *luxS* mutant. VE14089 is represented in grey bars with full outline and VE14089*ΔluxS* in white bars with a dashed outline. (A) Biofilm formation on polystyrene microtiter plates. The quantification of biofilm formation was assayed as a function of crystal violet stain (measured at 595 nm) retained by the biofilm biomass grown for 24 h. Data are mean of hexuplicate trials, and error bars indicate standard deviations. (B) VE14089 and VE14089*ΔluxS* adhesion to Caco-2 cells. The results are presented as per cent adhesion ± standard deviation. (C) Percentage (± standard deviation) survival of growing cells of *E. faecalis* VE14089 and VE14089*ΔluxS* at 2, 4 and 6 h of a challenge with 7 mM H_2_O_2._ One hundred per cent corresponds to the number of CFU before H_2_O_2_ treatment. All data are means (± standard deviation) of four independent experiments. (D) Time course of intracellular survival of VE14089 and VE14089*ΔluxS* strains within murine J774A.1 macrophages. Results correspond to the percent means ± standard deviations of intracellular Survival Index (SI) determined at 0, 4, 8 and 24 h post-killing of external bacteria, of five independent experiments. 0 h post-killing corresponds to 2 h post-infection, so time points mentioned in the figure correspond to 2, 6, 10 and 26 h post-infection.

### *luxS* mutation affects virulence in *Drosophila* septic injury model

*Drosophila* has recently been used successfully to test *E. faecalis* virulence factors (Teixeira *et al*., 2013). We thus decided to test the virulence of the *luxS* mutant in a septic injury *Drosophila* model. Flies injected with the *luxS* mutant strain showed delayed death when compared with flies injected with the wild-type *E. faecalis* strain (Fig. 5). After 24h post-injection, the *luxS* mutant allowed 60% survival of the injected flies, against the 20% survival of flies injected with the wild-type strain. Considering the results from the transcriptomic and metabolic *luxS* mutant profiles, the results shown in Fig. 5 suggest that the *luxS-*associated toxicity in the fly is likely related to one or several of the bacterial genes affected by the *luxS* deletion, leading to a lower cytotoxicity to the fly. This effect is significant and might be explored in future work in order to understand the lower toxicity to the host derived from the absence of the LuxS/AI-2 system.

**Figure 5.**
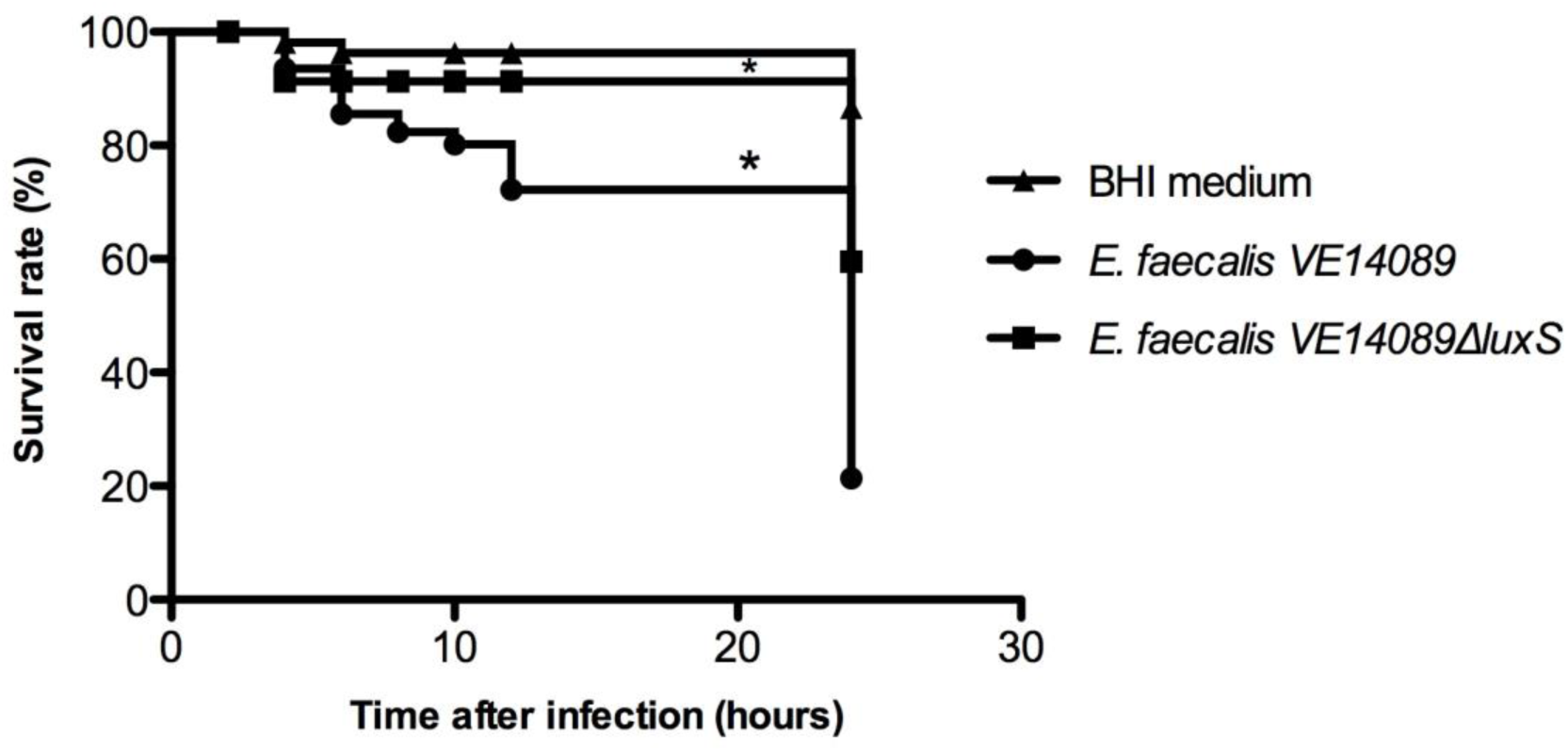
*Drosophila* survival rates upon infection with *E. faecalis* strains. 25 Oregon R (5-to 7-day-old) male adult flies, raised at 25°C, were infected, by septic injury onto the thorax with a thin needle, with VE14089 and VE14089*ΔluxS* strains. Data is representative of three independent experiments (75 flies per strain). Curves assigned with an * are significantly different (p< 0.0001) from the respective wild-type-infected curve, as determined by log-rank analysis.

## Discussion

The AI-2 activity in the culture supernatant of VE14089 was detectable from the mid-exponential phase onward, reaching maximum levels during late-exponential/early-stationary phase and then decreasing, while remaining higher than the basal levels detected during lag phase. AI-2 internalization and possible subsequent modification, through an ATP-binding-cassette transporter Lsr has been described for *Salmonella enterica serovar* Typhimurium (Taga *et al*., 2001), and a bioinformatics analysis showed that it is present also in other bacteria (Rezzonico & Duffy, 2008). This transport system has been hypothesized to be a mechanism to control AI-2 levels in the vicinity of a cell or to prevent AI-2 signaling by other bacterial species in its environment (Taga *et al*., 2003; Xavier & Bassler, 2005a; Xavier *et al*., 2007; Pereira *et al*., 2009). As an alternative to QS, after being released as a waste product, AI-2 may be reused subsequently as a metabolite (Winzer *et al*., 2002b), or as a borate scavenger (Coulthrust *et al*., 2002). When internalization occurs, the extracellular AI-2 concentration typically increases during exponential growth and begins to decline during the transition from exponential phase to the early stationary phase, by which time there are no detectable levels of AI-2 (Xavier & Bassler, 2005b; Azakami *et al*., 2006; De Keersmaecker *et al*., 2006; Han & Lu, 2009). The extracellular accumulation of AI-2 described for VE14089, as well as its level remaining considerably high, has also been described for other bacteria (Learman *et al*., 2009; Zhao *et al*., 2010) and may indicate the absence of AI-2 transport into the cells, which is in agreement with the genomic predictions previously made for *E. faecalis* (Rezzonico & Duffy, 2008).

Even if there is no internalization of the AI-2 molecule, a different mechanism of AI-2 detection in bacteria, like the one found in *V. harveyi*, can occur (Rezzonico & Duffy, 2008), where just the signal but not the AI-2 molecule is transduced inside the cell. This mechanism is triggered by the interaction of AI-2 with a two-component signal regulator pair (LuxP/LuxQ in *V. harveyi*), followed by a dephosphorylation cascade (Hardie & Heurlier, 2008). However, none of the published *E. faecalis* genomes contains potential homologues for the LuxP/LuxQ AI-2 signal transduction system found in *Vibrio* spp., which also happens for other bacteria.

Altogether, genome based predictions, the shape of the curve of AI-2 production during growth and the DPD inability to compensate for the transcriptional changes in the *luxS* mutant, all support the view of *E. faecalis* V583 as a bacterium blind to the interspecies communication molecule.

The *luxS* mutation did not significantly alter the expression of any of the genes involved in the AMC in VE14089, except for *luxS* (*ef1182*), the deleted gene. There were also no significant changes in expression of the putative methionine transporters or of the main pathways for methionine biosynthesis, from homoserine (or even the from the homoserine aspartate precursor) or serine precursors (data not shown). As predicted from Sri International Pathway Tools (v15.5) (Karp *et al*., 2010) V583 lacks the enzymes needed to convert homocysteine to methionine. Having no complete AMC for generation of methionine,, raises the question of whether the only function of LuxS in *E. faecalis* is as a QS molecule synthase, with a concomitant homocysteine generation, or whether LuxS in *E. faecalis* has another direct or indirect metabolic role, like maintaining the homeostasis of AMC metabolites and ensuring effective methylation (Heurlier *et al*., 2009). Accordingly, the *ruvAB* operon was highly down-regulated in the *luxS* mutant (Table 2). AMC provides methyl groups, among others, for DNA methylation. This, in turn, regulates important functions in cells, such as DNA repair, replication and transcription. Down-regulation of the *ruvAB* operon in the *luxS* mutant likely reflects lower methylation due to the absence of LuxS protein.

The absence of the LuxS enzyme may have other consequences besides changes in the methylation patterns of the cell. It is likely that deletion of the *luxS* gene in *E. faecalis* leads to different pools of AMC metabolites. For example, homocysteine is not produced, at least not from the AMC, and SRH likely accumulates from Pfs (EF2694) activity upon SAH. As the *luxS* mutant does not show any growth defect when compared to the wild-type strain, the metabolic rearrangements of the former, deduced from our transcriptomic analysis, likely reflect adaptation of the mutant cell to different activities of the AMC enzymes and the consequently different levels of AMC metabolites. It is possible that the increased utilization of BCAA, and other less preferred carbon sources, as energy sources observed in the *luxS* mutant is somehow related to regulatory networks activated by changes in the AMC metabolites. The demand of SAM is high during early and mid-exponential growth (Winzer *et al*., 2002a). This means that AMC enzymes are very active when cells are using preferred carbon sources, such as glucose. We can presume that, in the *luxS* mutant, the AMC is overall less active, which reflects upon the levels of metabolites such as homocysteine, SRH and SAM. It is possible that, under these circumstances, the cell interprets the lower active AMC as “we are in the stationary phase. Let’s derepress the utilization of less preferred carbon sources, such as BCAA”. Our bioinformatics analysis shows that several *cre* sites were found among the genes transcriptionally affected by deletion of the *luxS* gene. A role for catabolite repression in the activation of the BCAA degradation has been demonstrated in *E. faecalis* (Ward *et al*., 2000). Could AMC metabolite pools serve as intracellular signals for modulating catabolite repression? Recently, Redanz *et al.* (2012) demonstrated that *Streptococcus sangunis* biofilm defects, as well as most of the transcriptional changes of their *S. sanguinis luxS* mutant, could be restored to the wild type phenotype when an intact AMC was produced in the mutant by heterologous expression of *sahH* gene, coding for the enzyme that converts SAH into homocysteine. Their study provides evidence for a central role in metabolism of the AMC metabolites and presents an elegant strategy for future studies dedicated to understanding how AMC is regulated and influences metabolism and virulence in *E. faecalis*.

Interestingly, the *ef3194-ef3193* operon, which encodes a system made of genes *lrgA*, encoding a putative murein hydrolase regulator holin-like protein, and *lrgB*, encoding an antiholin-like protein, was highly up-regulated in the *luxS* mutant strain. Although the role of these genes in *E. faecalis* is not yet elucidated, it is known that they are induced by the LytRS regulatory system, contribute to *E. faecalis* infection in *Drosophila* (Teixeira *et al*., 2013) and are induced during growth in blood (Vebo *et al*., 2009). *lrgAB* is thought to be induced by the LytRS system (Teixeira *et al*., 2013), which, in *S. aureus* (Bayles, 2007), senses decreases in membrane potential caused by proton motive force. Despite a role in autolysis regulation proposed in other species (Lui *et al*., 2011), *lrgAB* expression is also modulated by the metabolic activity of cells and may have been hugely up-regulated in the *E. faecalis luxS* mutant in response to the massive metabolic reorganization due to the absence of the LuxS protein.

Microarray results show that known virulence factors of *E. faecalis* were not transcriptionally affected by the absence of an active LuxS protein or AI-2. This finding is in accordance with results observed in assays associated with *E. faecalis* pathogenic behavior. However, and as previously reported for other bacteria, *luxS* deletion leads to major metabolic reorganization in *E. faecalis* cells, involving both sugar transport and metabolism and also of branched-chain amino acids. Biolog plate screening showed that *luxS* deletion induced an increased and faster growth on galactose. It is possible that the delayed death of flies by the *luxS* mutant strain observed during *Drosophila* infection is related to different metabolism of the wild-type and *luxS* mutant strains. This result warrants, however, further investigation as any attempt to decrease *E. faecalis* infectivity during sepsis is of huge importance and urgency. In particular, metabolism may have greater impact on virulence of a strain when tested in polymicrobial habitats, as recently shown by Ramsey *et al*. (2011). Further studies should clarify if the metabolic change induced by *luxS* deletion and AI-2 absence has an impact on *E. faecalis* virulence in polymicrobial infection and/or colonization models. However, we cannot exclude the possibility that the observed decreased virulence associated to *luxS* mutation might be related to the absence of AI-2 produced by this strain. Although there are no published works reporting AI-2 effects on the host itself, other autoinducer molecules produced by bacteria are known to affect, for example, the immune system activation of the host (Hughes & Sperandio, 2008).

Altogether, this work evidences the importance of LuxS in *E. faecalis*. Our findings point to a role of this protein at the metabolic level. Moreover, impaired LuxS activity led to attenuated virulence in *Drosophila*. However, none of the other phenotypes tested, namely adhesion to Coco-2 cells, survival inside macrophages, resistance to oxidative stress and biofilm formation, was affected by *luxS* deletion. Recently, Shao *et al*. (2012) claimed that AI-2 promotes biofilm formation by V583 strain. Despite being statistically relevant, they report that the increase in biofilm by AI-2 addition was very small (OD increased from 0.25 to 0.3 with 0.20 µM AI-2). Using a proteomic approach they propose that AI-2 signal regulates some metabolic features in *E. faecalis*. Despite this conclusion, our findings did not overlap with theirs. Differences between ours and Shao *et al*. (2012) results are most likely due to differences in experimental design. In their study the V583 strain was carrying a fully functional LuxS protein and therefore the role of the *luxS* gene was not truly tested. Moreover, we used DPD whereas they used AI-2 prepared *in vitro* from SAH which was enzymatically degraded by purified LuxS and Pfs enzymes. Therefore, the two studies should not be compared.

LuxS activity in *E. faecalis* was shown to lead to production and extracellular release of AI-2. Despite *E. faecalis* inability to sense this molecule, at least under the conditions tested, it may be considered as an indicator of *E. faecalis* metabolic activity by other bacteria able to sense AI-2 and respond to it. *E. faecalis* colonizes mainly environments where multispecies communities are present, such as those in food, human gut and soil, and often appears in polymicrobial infections. The fact that *E. faecalis* is unable to respond to AI-2, either self or from others, does not mean that the self-produced AI-2 does not have implications in the environmental niches it occupies. Czárán and Hoekstra have proposed a model for communication, cooperation and cheating in multispecies communities (Czaran & Hoekstra, 2009). According to their model, eight different behaviors are possible when quorum-sensing is considered based on presence/absence of three genetic loci: cooperation (production of a public good); production of the quorum signal molecule; and response to the quorum molecule. According to this model, *E. faecalis* would be considered a “liar” as it does not cooperate: it produces the quorum signal, do not respond to it and no common goods appear to be produced. *E. faecalis* may benefit from others by being non-cooperative but inducing others to change the environment for its own benefit. This behavior could also be called “cheating”, so defined when a cell does not cooperate, but benefits from public goods produced by cooperating bacteria (West *et al*., 2006). Both liar and cheater behaviors are based upon the assumption that AI-2 functions as a signaling molecule for communication between *E. faecalis* and other species, which might not hold true for all the species that co-share environments with *E. faecalis*. According to Diggle et al. (2007), the fact that AI-2 produced by one species can influence gene expression in another species does not mean that we can generalize it as a signaling molecule in all interspecies interactions. In some microbial interactions involving interspecies interactions, AI-2 could fit into the category of a cue or of a coercive molecule. The first is applicable to cases when the production of a substance by individual A has not evolved because of its effect on individual B (Maynard-Smith & Harper, 2003; Diggle, 2010); the second labels a case when the production of a substance by individual A forces a costly response from individual B (Diggle et al., 2007; Maynard-Smith & Harper, 2003; Keller & Surette, 2006; Diggle, 2010). It would be interesting to know the exact nature of *E. faecalis* social behavior associated to AI-2 production, if it is a liar, cheater or coercive interaction. Understanding if and how *E. faecalis* benefits from public goods produced by bacteria responding to AI-2 could explain the opportunistic nature of *E. faecalis* and we could benefit from future exploitation of this knowledge by developing new antimicrobial strategies.

## Acknowledgements

We are grateful to Catarina Pereira and Karina Xavier for their help and assistance in performing AI-2 quantification, done in Karina Xavier’s Laboratory (ITQB/IGC). The authors acknowledge funding by Fundação para a Ciência e Tecnologia for project grant PTDC/CVT/67270/2006, co-financed through FEDER, and PEst-OE/EQB/LA0004/2011, and for grant SFRH/BD/18757/2004 attributed to Frédéric Gaspar.

